# Root density sensing allows pro-active modulation of shoot growth to avoid future resource limitation

**DOI:** 10.1101/539726

**Authors:** Cara D. Wheeldon, Catriona H. Walker, Tom Bennett

## Abstract

Plants use environmental cues to determine their optimal root and shoot growth. It is well known to gardeners and horticulturists alike that soil volume – most commonly in the form of pot size – strongly restricts plant growth, but the mechanisms by which this effect occurs remain unclear. Here, we show that shoot growth scales directly with soil volume, independently of the nutritional content of the soil, and that plants can become ‘volume restricted’ even in the presence of abundant resources. We show that plants can detect their soil volume as early as 3 weeks post germination, and that shoot growth restriction therefore constitutes a pro-active ‘decision’ by the plant to avoid resource limitation later in the life cycle. Shoot growth restriction is not directly linked to root growth restriction, and does not occur in response to the mechanical detection of the pot walls. Rather, we show that plants detect their soil volume by detecting the density of roots in the proximity of their root system. As such, volume restriction may be intimately connected with the mechanism by which plants sense and respond to the roots of other plants in the rhizosphere. Our work demonstrates the remarkable ability of plants to make pro-active decisions about their growth to ensure they can complete their cycle, and has important implications for agricultural practise regarding both nutrient use efficiency and yield.

## INTRODUCTION

The growth and development of a typical plant is strongly intertwined with its immediate environment. Plants need to acquire resources (light, water, mineral nutrients) in order to grow, and as such, the majority of the plant body consists of tissues specialised for acquiring those resources; roots, leaves and homologous structures. Thus, while on the one hand plant growth can be limited by the scarcity of available resources in the environment, it is also true that there is little benefit in increasing the amount of resource-harvesting structures if there are no resources to harvest. The relationship between plant growth and resource availability is therefore more complex than often portrayed. While it is certainly true that plant growth can be absolutely limited by lack of resources, it is also the case that plants can pro-actively ‘choose’ not to invest in more growth. Indeed, where sufficient information is present in the environment to allow plants to predict future resource scarcity, it is a far more useful strategy to pro-actively limit growth to avoid resource limitations, than to grow until resources are exhausted (Walker & Bennett, 2018). We can characterise changes in plant development as ‘decisions’ if they are taken in the absence of resource limitations, and ‘responses’ if they occur due to resource limitations. There are plenty of reasons why plants should make pro-active decisions about their growth. Life-cycle constraints are a particularly good example; if a plant only has available resources to reach a certain biomass, it should not reach that size while still vegetative, otherwise it will be unable to complete its life cycle. Thus, the plant must actively constrain its early growth (even though it has available resources) to leave resources available for reproduction in the future.

In vascular plants, root and shoot systems fulfil completely opposite functions in resource harvesting, and as such are mutually interdependent. To grow optimally for given conditions, plants must therefore make decisions about investment in shoot growth with respect to the availability of resources in the root system, and vice versa. The ability of plants to make such complex and coordinated decisions about their growth and development is made possible by extensive long-distance signalling across the plant body. This signalling occurs through the production and perception of a small handful of ‘phytohormones’, of which auxin, cytokinin (CK), strigolactone (SL) and gibberellin (GA) are particularly relevant for coordinating growth over long-distances. Since these molecules each regulate a plethora of developmental processes, they are perhaps best thought of as ‘informational’ rather than instructive; they relay information about environmental conditions from one part of the plant to the rest (Bennett & Leyser, 2014). Each organ can use this information to respond appropriately to conditions. Alongside these phytohormones, a range of small peptide signals with more specific roles also act in long-distance signalling (reviewed in Matsubayashi, 2014). In the context of root-shoot coordination, auxin acts as major shoot-root signal, while SL and CK are major root-shoot signals. For instance, a relatively large body of work in recent years has demonstrated how these hormones, and their interactions with each other, allow plants to regulate their shoot branching with respect to nitrate (N) and phosphate (P) availability in the soil (Umehara et al, 2010; Kohlen et al, 2011; de Jong et al, 2014; Muller et al, 2015; Waldie et al, 2018).

Given their status as the major mineral nutrients, the strong influence of external N and P availability on plant growth is unsurprising. The availability of water could also be used to make pro-active decisions regarding shoot growth, since producing a larger shoot will create a higher peak demand for water. It is certainly clear that lack of water has strong effects on plant growth, but these are usually stress responses (e.g. Osakabe et al, 2014) and where tested, water availability does not seem to strongly influence decision-making and shoot architecture (Shemesh et al, 2012). Ultimately, water availability may be too unpredictable to use as a reliable cue for future growth. Besides its chemical constituents, there are other physical and biological components of the rhizosphere that plants could use to predict the likelihood of resource availability in the future. For instance, soil depth might act as a proxy for likely water availability at the height of summer. The influence of soil volume - or often more specifically, pot size - on shoot growth has previously been described by a range of studies. Reduction in shoot growth is a commonly reported finding across a range of species such as bean, cotton and tomato, when available rooting volume is reduced (Carmi & Heuer, 1981; Yong et al, 2010; Bar-Tal et al, 1995; Poorter et al, 2012). This shoot growth reduction does not appear to be correlated with a reduction in photosynthetic capacity, with studies reporting an increase, decrease or no change, in reduced pot sizes (Kharkina et al, 1999; Krizek et al, 1985; Shi et al, 2008; Yong et al, 2010). These effects are also not caused by water deficit (Krizek et al, 1985; Ismail & Davies, 1998) and indeed, still occur in hydroponic conditions (e.g. Ternesi et al, 1994; Shi et al, 2008). Furthermore, these effects do not appear to correlate with the availability of nutrients in the container (Bar-Tal et al, 1995), a view supported by a recent meta-analysis of 65 different pot size studies (Poorter et al, 2012).

Based on these findings, we hypothesised that the plants use soil volume as a predictive factor for future nutrient and water availability, and that responses to soil volume would therefore constitute pro-active decisions, rather than simple responses. In this study, we have tested these ideas, and attempted to understand how plants might sense soil volume. We show that shoot growth scales with soil volume in a linear fashion up to certain limit, at which plants reach a maximum inherent size. This effect is non-nutritional, and soil volume and nutrient concentration can simultaneously modulate growth. We show that volume restriction effects on shoot growth can begin as early as 3 weeks after germination, and do not co-occur with root growth inhibition. Rather, volume restriction is a function of the effective root density of the root system. As such, the detection and effects of soil volume on shoot growth may be closely related, if not identical, to the mechanism by which plants detect and respond to the presence of other plants in the rhizosphere.

## RESULTS

### Pot size causes a non-nutritional limitation on shoot growth, seed set and yield

To understand how available soil volume influences shoot growth, we grew spring wheat (*Triticum aestivium*; two varieties, Mulika and Willow), spring barley (*Hordeum vulgarum*; two varieties, Charon and Propino) and spring oilseed rape (OSR; *Brassica napus*; one variety; Heros) in pots containing either 100ml, 500ml or 2000ml of soil, under well-illuminated and well-watered conditions. We also grew Arabidopsis (*Arabidopsis thaliana*; Col-0 wild-type) in pots containing either 50ml, 100ml of 500ml of soil, in similar conditions. We observed that in each species, the size of the shoot system was clearly proportional to the size of the pot (Figure 1A, B). We measured shoot size using the number of shoot branches as a proxy for shoot system size in Arabidopsis and OSR, and the peak number of tillers (the equivalent parameter) in wheat and barley. These results clearly illustrate the effect of soil volume on shoot growth; there is a linear, direct proportionality between pot size and branch number (Figure 1C, G; Figure S1A, C). The same linear relationship with respect to soil volume is observed if we instead measure shoot biomass (fresh biomass for OSR, dry biomass for Arabidopsis, dry straw biomass for wheat and barley) as a proxy for shoot system size (Figure 1D, H; Figure S1B, D). Thus, as previously demonstrated in many species (reviewed in Poorter et al, 2012), soil volume clearly acts as a direct constraint on shoot system size.

**Figure 1:**
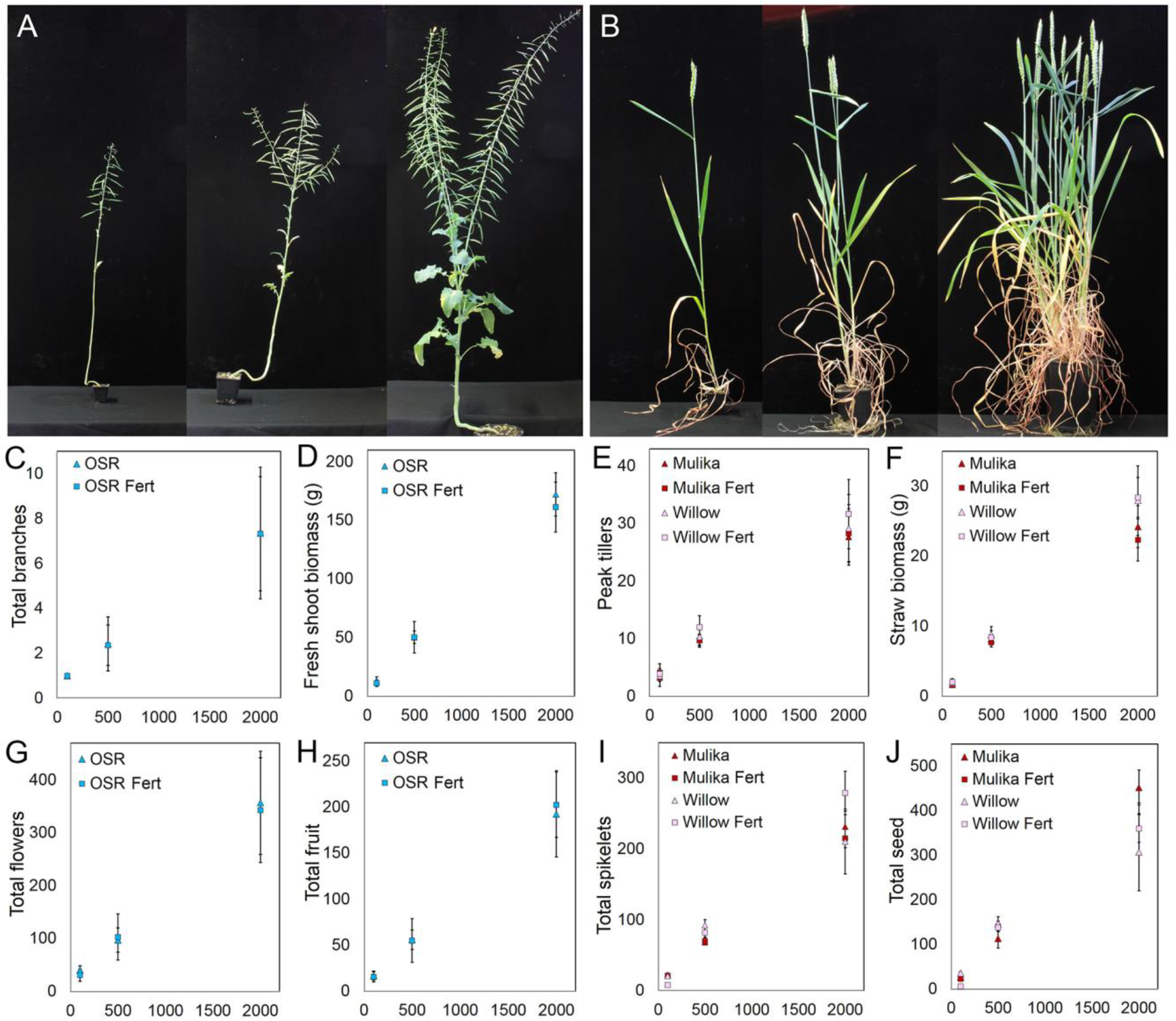
Soil volume directly influences plant growth. **A, B)** Final plant size in spring oilseed rape plants (A) and spring wheat plants (B) grown in 100, 500 and 2000ml of soil. **C,D,G,H)** Graphs showing the relationship between soil volume and mean total branch number (C), mean fresh shoot biomass in grams (D), mean total flowers (G) and mean total fruit (H) in spring oilseed rape grown in 100, 500 and 2000ml of soil, with (‘Fert’) or without additional fertiliser. Error bars indicate s.e.m, n=6-12. **E,F,I,J)** Graphs showing the relationship between soil volume and mean peak tiller number (E), mean dry straw biomass in grams (F), mean total spikelets (I) and mean total seed (J) in two varieties of spring wheat (Mulika and Willow) in 100, 500 and 2000ml of soil, with (‘Fert’) or without additional fertiliser. Error bars indicate s.e.m, n=6-12.

We also investigated the effect of soil volume on reproduction in these species, especially since seed yield is a much more important parameter in many crops than the size of the shoot system *per se*. For OSR, we measured the number of flowers and fruit produced by plants in different pot sizes. As anticipated, there was a linear relationship between flower/fruit number and soil volume (Figure 1E, F). In wheat, we measured a number of different parameters for reproduction including the number and biomass of ears, the number of spikelets and the number and biomass of seeds produced (Table S1). For each parameter, we observed the same linear relationship with pot size (Figure 1I, J; Table S1). Thus, for any shoot growth parameter we measured, whether vegetative or reproductive, we observed that soil volume strongly influenced the growth of plants in a linear manner, over the tested range.

One possible explanation for this effect is that, since different soil volumes will contain different quantities of essential mineral nutrients such as nitrate (N) and phosphate (P), the growth of the plants is limited by the availability of nutrients. We reasoned that if this was the case, adding additional fertiliser to the pots would alleviate the growth restriction relative to control plants. Thus, in all the experiments described above, we treated half the plants in each pot size with additional fertiliser (10ml of standard nutrient solution per plant per treatment, weekly), at rates that are comparable to those rates commonly used in agricultural practise (i.e. ∼80-200 kg/ha of N). However, we did not observe any significant increase in shoot system size in any of the plants treated with additional fertiliser relative to control plants, whether judged by biomass or branching, seed-set or yield (Figure 1C-J; Figure S1A-D; Table S1). Thus, consistent with indications from various previous studies (Hameed et al, 1987; Bar-Tal & Pressman, 1996; Poorter et al, 2012) we conclude that the effect of soil volume on plant growth is not a nutritional effect, but rather a separate stimulus that acts independently of nutrient levels.

### Complex interplay of nutrient availability and soil volume on shoot growth

In agricultural systems, the availability of mineral nutrients, particularly nitrate (N) is often a strongly limiting factor for crop growth. We therefore found it striking that there was no effect on plant growth of additional nutrients under volume-restricted conditions (Figure 1). This suggests that for a given volume of soil, there may be a maximum level of nutrient uptake by plants, and that the compost used in our experiments exceeds this level of nutrients even without additional fertiliser. However, we reasoned that at sufficiently low nutrient levels, the effect of additional nutrients would be more obvious, and the effect of pot size should therefore be reduced. To test this idea, we used a system where we could precisely control nutrient inputs, by growing plants in a nutrient-poor medium of sand/vermiculite or sand/perlite. Using this system, we grew wheat (Mulika) in two pot sizes (100ml and 500ml) supplemented by weekly fertiliser with either a ‘low’ nitrate (N) concentration (7.5µmol of N/week) or a ‘standard’ N concentration (75µmol of N/week). All plants received the ‘standard’ concentration of other nutrients, and different pot sizes received the same total amount of fertiliser each week. In 2/3 independent experiments, we observed no statistically significant effect of pot size on shoot biomass under low N treatment, with biomass being largely determined by nitrate availability (Figure 2A). However, in these experiments, there was a clear effect of pot size on shoot biomass under standard N conditions, with plants grown on 500ml of soil accumulating more biomass than those on 100ml; in both pot sizes, plants accumulated more biomass than the equivalent low N-treated plants (Figure 2A).

**Figure 2:**
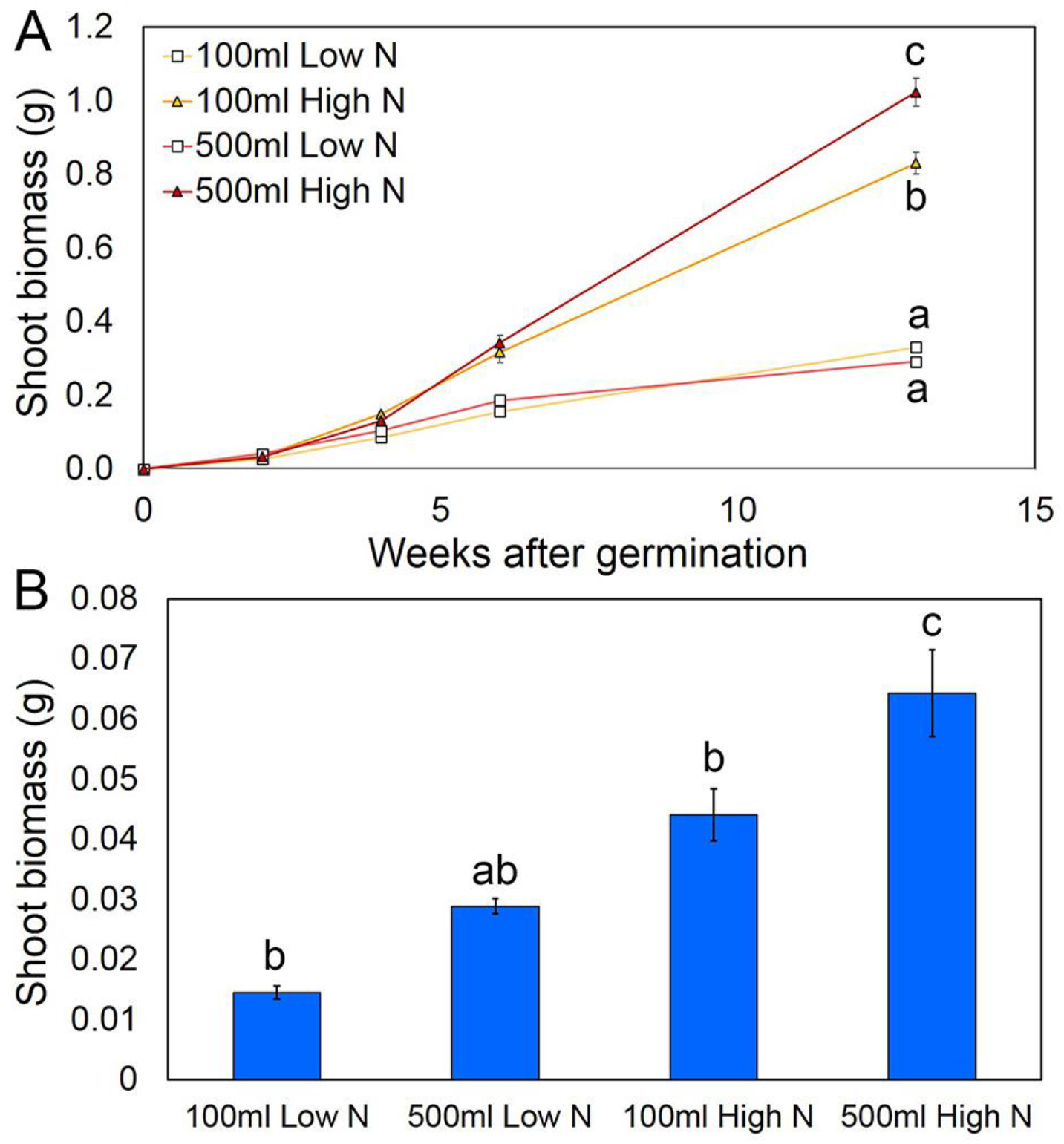
Soil volume and nutrient availability both regulate shoot growth. **A)** Graph showing mean dry shoot biomass over a 13-week period in spring wheat plants grown on a sand/vermiculite mix, in two pot sizes (100 and 500ml) and supplemented with fertiliser containing either standard nitrate concentration (75µmol/week) or a low nitrate concentration (7.5µmol/week). Data points are means ± s.e.m. n=4 plants for weeks 2, 4, and 6, n=8 for week 13. **B)** Graph showing mean final dry shoot biomass in wild-type (Col-0) Arabidopsis grown on a sand/vermiculite mix, in two pot sizes (100 and 500ml) and supplemented with fertiliser containing either standard nitrate concentration (75µmol/week) or a low nitrate concentration (7.5µmol/week). Data points are means ± s.e.m. n=10. Bars with the same letter are not statistically different from each other (ANOVA+Tukey HSD).

We also performed similar experiments using Arabidopsis, grown in 100 and 500ml of sand/perlite mix, supplemented with low N or standard N fertiliser (7.5 or 75µmol of N/week). Despite its diminutive size, we observed a stronger effect of pot size in Arabidopsis than in wheat. Even under low N treatment, plants grown in larger pots had significantly greater shoot biomass (Figure 2B), than in smaller pots. This would be consistent with Arabidopsis having a much lower demand for N than wheat, and therefore becoming volume restricted at lower N concentrations. We observed the same trend under standard N treatment, with plants in larger pots having increased shoot biomass relative to those in smaller pots (Figure 2B). Plants grown in standard N conditions were larger than those grown in the same pot size in low N conditions (Figure 2B). Thus, being volume restricted did not prevent plants grown in 100ml pots from responding to increased N availability in these experiments.

Collectively our results from Arabidopsis and wheat suggest that at very low nutrient conditions, plant growth is entirely limited by nutrient availability, while at very high nutrient conditions growth is entirely limited by soil volume. In between these extremes, there are a range of nutrient levels at which growth is sensitive to both nutrient levels and soil volume. The separable effects of pot size and nutrient availability shown here also further emphasise that the effect of volume restriction on shoot growth is non-nutritional.

### The effect of soil volume on shoot growth is saturable

Having demonstrated that for a given pot volume, the effect of nutrients on growth is saturable, we next questioned whether the effect of soil volume on shoot system size was continuous, or whether there was a point at which plants were no longer ‘volume restricted’. To test this idea, we grew Arabidopsis in a standard volume of soil (100ml) and three over-sized volumes (500, 1000 and 2000ml). We observed a large increase in total branch number between 100ml and 500ml, as expected (Figure 3A). However, between 500ml and 1000ml there was only a marginal and statistically insignificant increase in branch number despite a doubling in soil volume; and similarly between 1000ml and 2000ml (Figure 3A). Thus, Arabidopsis growth in relation to pot size seems to plateau in the range 500-1000ml, above which plants are essentially ‘unrestricted’. This is also mirrored in the small, non-linear increase in biomass between plants grown in 500 and 2000ml of soil (Figure 3B). In comparison, the relationship between pot size and shoot size is still linear between 500ml and 2000ml for wheat, barley and OSR. Thus, the point at which a plant ‘escapes’ from volume restriction seems to be a function of the inherent size of the species in question.

**Figure 3:**
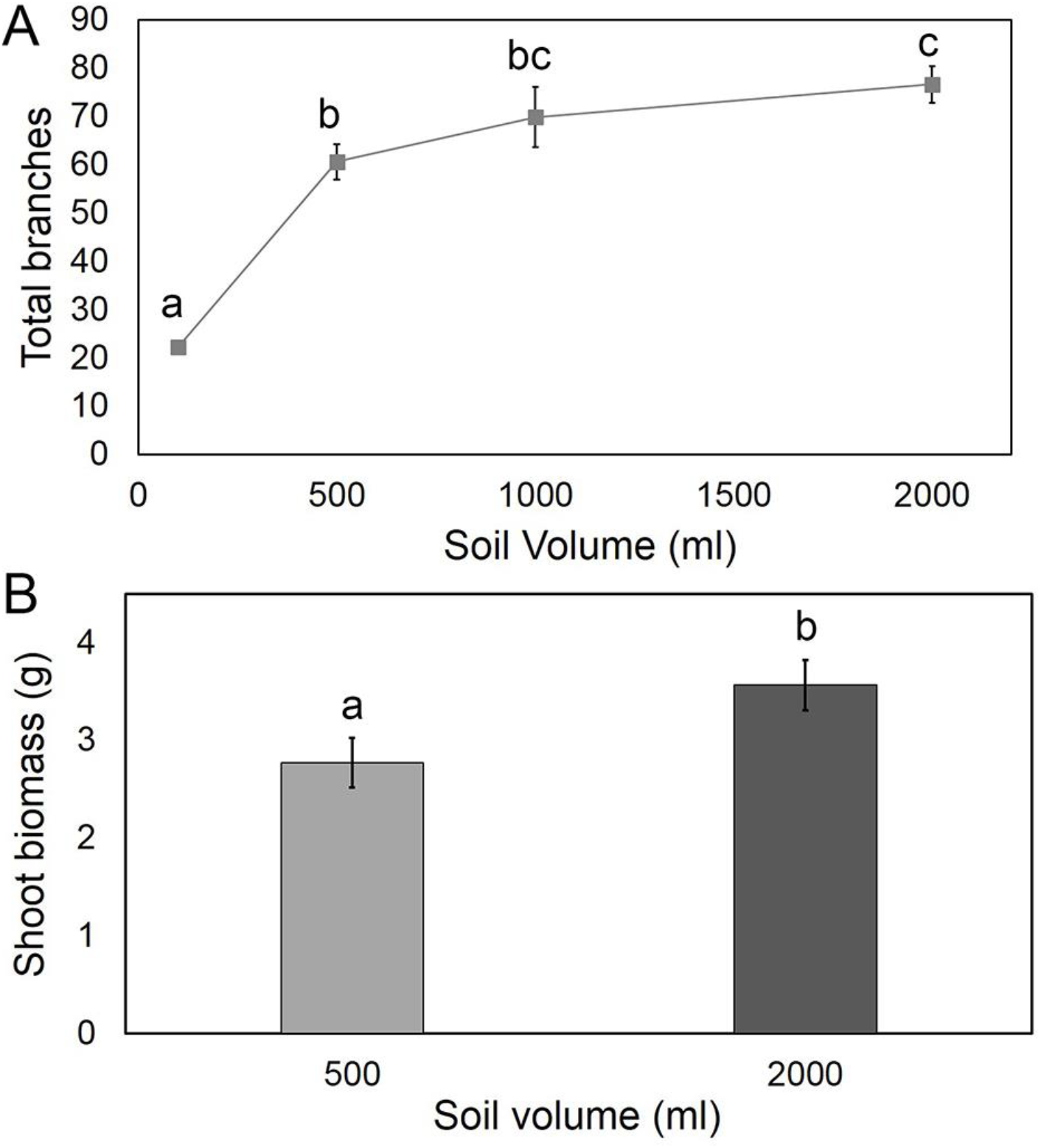
Shoot growth does not scale indefinitely with soil volume. **A)** Graph showing mean total branch number in wild-type (Col-0) Arabidopsis grown on compost, in four pot sizes (100, 500, 1000 and 2000ml). Data points are means ± s.e.m. n=8-12. Bars with the same letter are not statistically different from each other (ANOVA+Tukey HSD). **B)** Graph showing mean final dry shoot biomass in wild-type (Col-0) Arabidopsis grown on compost, in two pot sizes (500 and 2000ml). Data points are means ± s.e.m., n=10. Bars with the same letter are not statistically different from each other (T-test, p<0.05).

### Shoot responses to volume restriction are pro-active

We wanted to understand whether the effects of volume restriction on shoot growth constitute a reactive response to stresses caused by the limited volume, or whether they constitute a proactive decision, informed by the environmental conditions, that avoids such stresses occurring. The growth habit of wheat and barley provide an excellent method for testing this, because they initiate basal branches (tillers) from very early in the life-cycle. We therefore tracked tiller number in spring wheat and spring barley grown in 100ml, 500ml and 2000ml pots over 16 weeks (Figure 4A, B). In all plants, tillering began after 3 weeks of growth, and almost immediately, the plants in different pot sizes began to diverge in terms of the rate of tiller production. Thereafter all plants continued to initiate tillers, before reaching peak tiller numbers until 6-7 (100ml) or 8 weeks after germination (500ml and 2000ml). Later in development a proportion of tillers underwent senescence, until by 14 weeks after germination, all remaining tillers have initiated an ear. The point at which the different pot sizes diverge (3 weeks) must represent the point at which the plants have detected the limitation in their soil volume, and begin modulating their shoot architecture with respect to it. However, this is not the point at which plants stop tillering, even in the smallest soil volume; they continue to initiate new tillers for at least another 2-3 weeks after this point. Furthermore, even when tillering stops, the plants clearly continue to grow (in terms of biomass; Figure 2A) and develop (in terms of life-cycle progression; Figure 1B) until the normal end of their life. In separate experiments, we were able to grow wheat plants for at least 3 weeks in un-supplemented vermiculite, with the plant solely utilising the nutrients stored in the wheat grain (Figure 4D, E). Moreover, plants provided with weekly additional fertiliser (10ml of standard nutrient solution) do not have different growth curves relative to untreated plants (Figure 4C). Thus, again, our data conclusively demonstrate that the divergence in tillering between pot sizes cannot be a response to nutrient levels in the soil.

**Figure 4:**
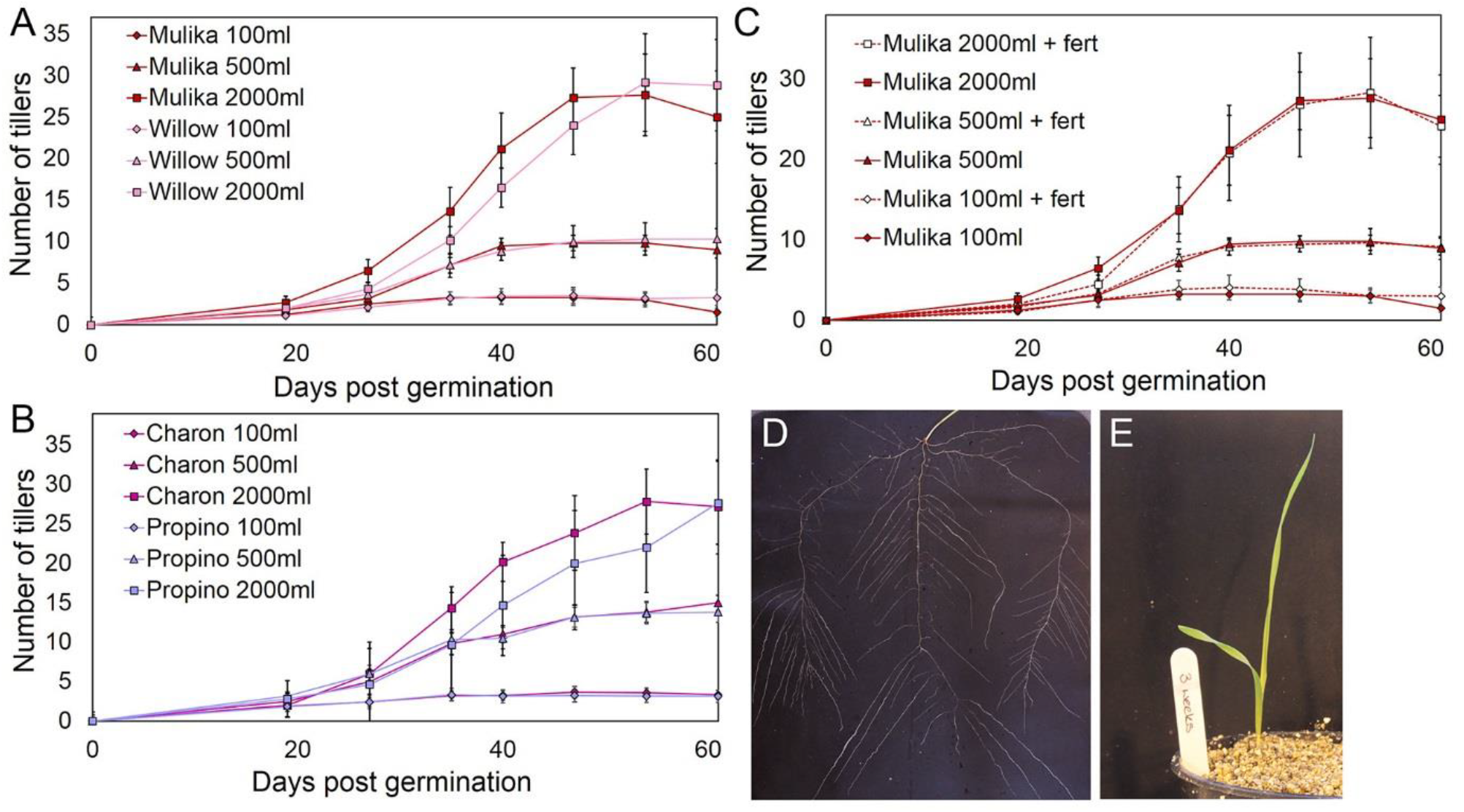
Plants respond early in the life-cycle to soil volume. **A)** Graph showing mean tiller number in the 63 days after germination in two varieties of spring wheat (Mulika and Willow) in 100, 500 and 2000ml of soil. Error bars indicate s.e.m, n=6-12. **B)**Graph showing mean tiller number in the 63 days after germination in two varieties of spring barley (Propino and Charon) in 100, 500 and 2000ml of soil. Error bars indicate s.e.m, n=6-12. **C)**Graph showing mean tiller number in the 63 days after germination in spring wheat (Mulika) in 100, 500 and 2000ml of soil, with additional fertiliser (‘+ fert’) or without additional fertiliser (same lines as shown in A). Error bars indicate s.e.m, n=6-12. **D, E)** Root (D) and shoot (E) system growth in wheat grown in ultra-low nutrient substrate, 3 weeks post germination.

The clear implication of these results is that the plants are able to detect their soil volume very early in their life-cycle, and pro-actively adjust the scale of their shoot system (in architectural terms) in response. As a result of these architectural changes they also adjust their overall rate of growth, but plants continue accumulating biomass throughout their life even in the most limiting conditions (e.g. Figure 2A). These pro-active changes in architecture thus allow plants to successfully complete their life cycle despite the limitations of their environment, rather than simply growing at the same rate in all conditions until they are run out of resources and are unable to grow further. Plants do not wait for resource limitation to occur, but rather use environmental information to ‘anticipate’ the possibility of resource limitation and modify their shoot architecture to avoid becoming resource-limited.

### Changes in shoot growth are not directly correlated with root growth

Our results demonstrate that wheat plants can detect their soil volume early in the life-cycle and respond pro-actively to limit their shoot branching. We wanted to understand how root growth responds to soil volume and nutrient level, and whether changes in root growth might directly influence shoot growth. We thus measured root biomass in wheat plants grown on sand/vermiculite or sand/perlite, as described above. As expected, for low N treatments, there was generally little effect of pot size on root growth, with biomass being largely determined by nitrate availability (Figure 5A, B). The effect of pot size on root biomass accumulation at standard N concentration was more variable, but in 2/3 experiments, root biomass was not significantly higher in the larger pots (Figure 5A). In one experiment, we tracked root biomass accumulation across the duration of the experiment. In all treatments, root growth was at its highest between 2-6 weeks post-germination, the window in which tillering normally occurs (Figure 5). However, no tillers were formed in any plants in this experiment; inhibition of tillering is thus not correlated with inhibition of root growth. Conversely, the differences in final shoot biomass between pot sizes in the standard N treatment arise after week 6, once root growth is largely stalled, and in the absence of any difference in root growth between pot sizes (compare Figure 5 and Figure 2A). Changes in shoot growth in this system therefore do not directly arise as a result of altered root growth.

**Figure 5:**
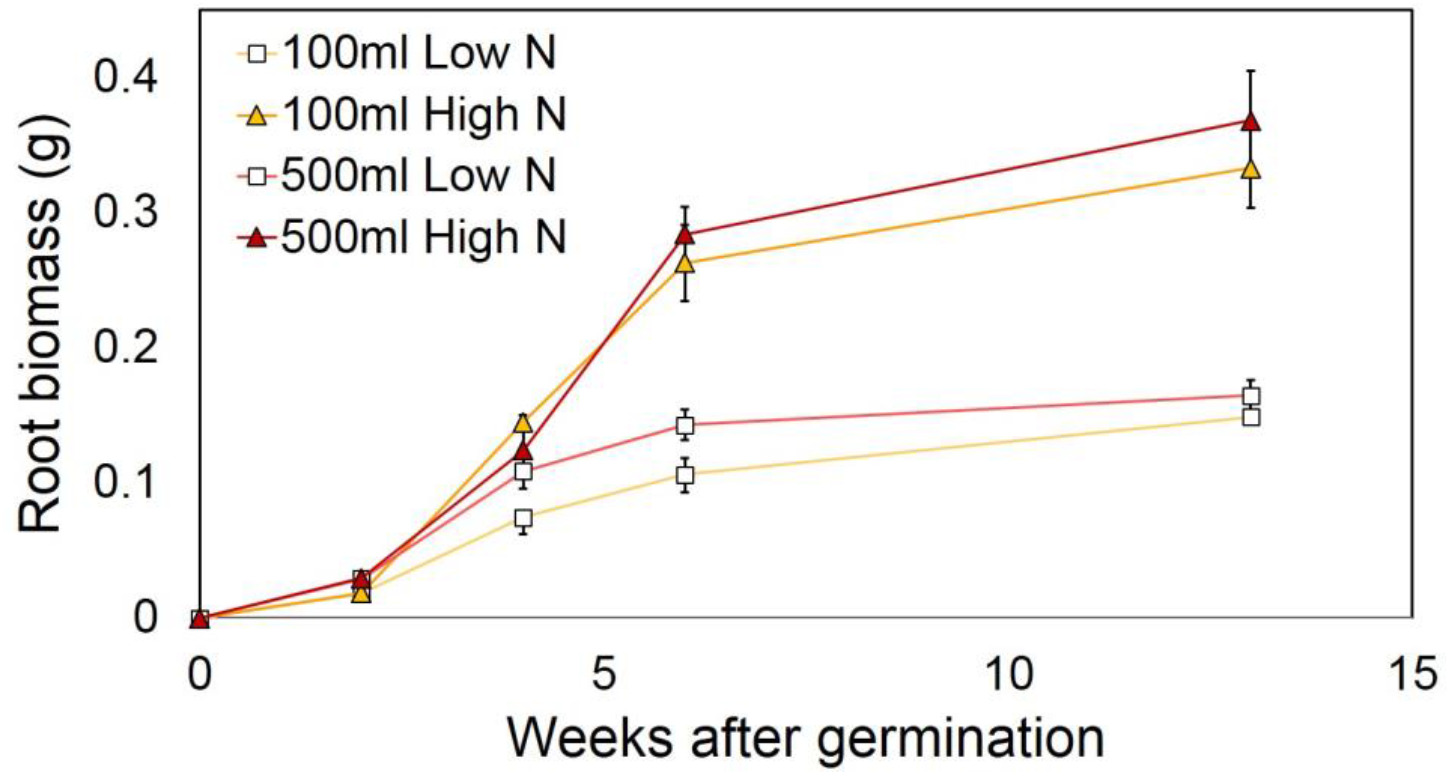
Root growth does not directly determine shoot growth in restricted plants. Graph showing mean dry root biomass over a 13-week period in spring wheat (Mulika) plants grown on a sand/vermiculite mix, in two pot sizes (100 and 500ml) and supplemented with fertiliser containing either standard nitrate concentration (75µmol/week) or a low nitrate concentration (7.5µmol/week). Data points are means ± s.e.m. n=4 plants for weeks 2, 4, and 6, n=8 for week 13.

### Volume restriction is unlikely to occur through mechanical effects

Our data suggest that volume restriction is a pro-active response, which likely occurs during active root growth, and not as a consequence of root growth inhibition. We hypothesised that two alternative mechanisms might explain how plants detect volume restriction. Firstly, we postulated that volume restriction might be primarily detected through mechanical interaction of root growth by the pot walls, or secondly, that is detected as a function of root density. To test these ideas, we grew wheat plants (Mulika) in a soil-vermiculite mix in clear-walled containers of two different sizes (300ml and 1100ml), allowing us to inspect the progression of root growth in relation to shoot growth. We observed that, within the first week post germination, roots had already collided with the walls of both pot sizes (Figure 6A), and deflected their growth along the wall. As expected, the plants in both pot sizes began to tiller after 2 weeks, and the initiation of tillers was identical between pot sizes until 4 weeks post germination, after which they diverged, with tiller formation peaking in smaller pots before 6 weeks post germination (Figure 6B). At this point, there were large numbers of roots in contact with the pot walls in both pot sizes, and no obvious difference between the pot sizes in this regard (Figure 6C-F). However, the density of roots was clearly much higher in the smaller pots, especially along the bottom wall of the pot (Figure 6E). These data suggested that mechanical impedance was not correlated with changes in shoot growth, but that root density might be.

**Figure 6:**
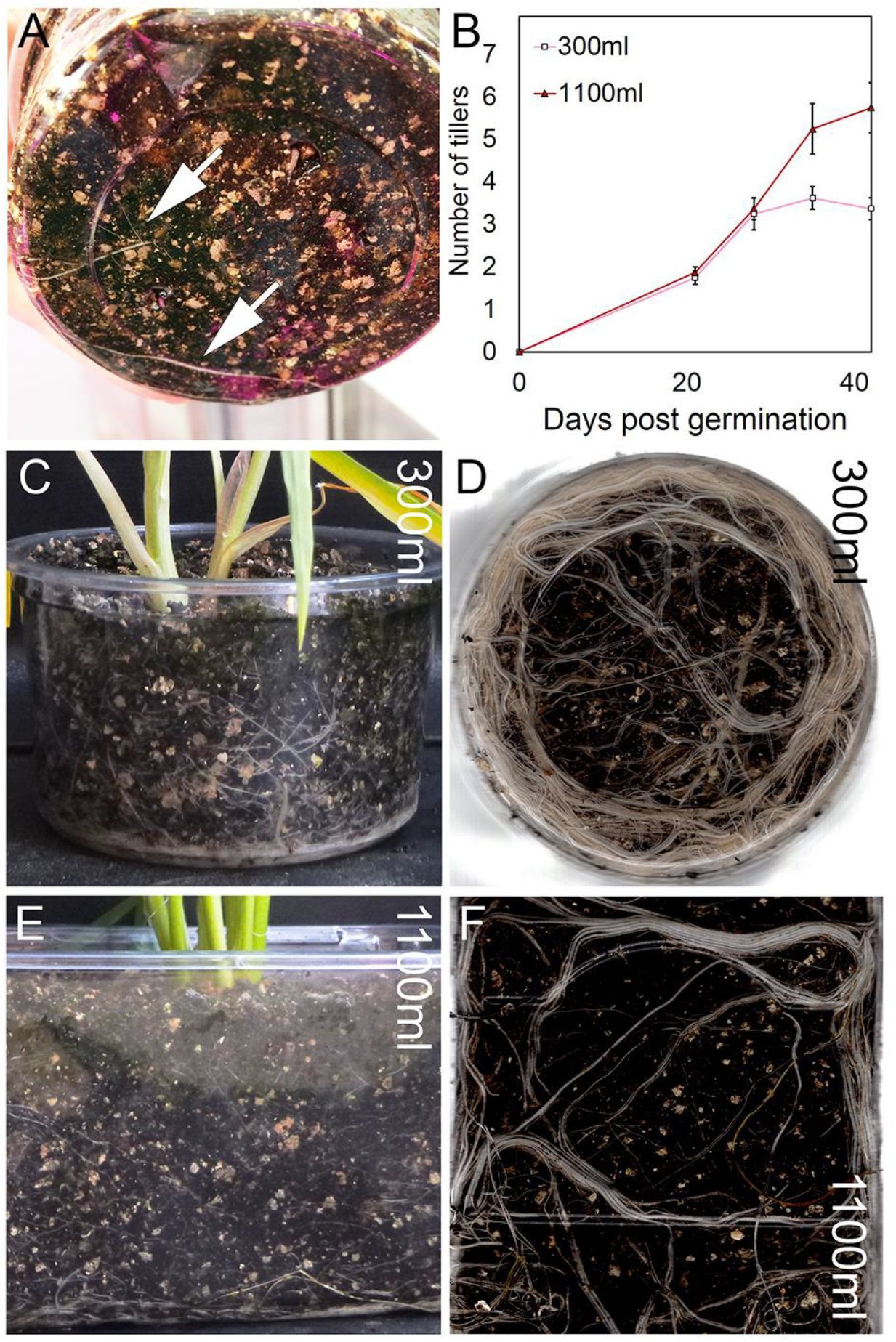
Shoot growth inhibition does not correlate with mechanical stimulus. **A)** Root development in spring wheat grown in clear sided 300ml pots, 1 week after germination. **B)**Graph showing mean tiller number in the 42 days after germination in spring wheat (Mulika and Willow) grown in 300 and 1100ml of soil/vermiculite mix. Error bars indicate s.e.m, n=8. **C-F**) Root development visualised through the pot walls of clear sided 300 (C,D) and 1100ml (E,F) pots, 42 days after germination.

### Volume restriction is a function of plant root system density

To test this idea more formally, we performed an experiment in which we grew 3 groups of wheat plants (Mulika) in 100ml pots for 4 weeks, alongside 1 group of plants in 500ml pots. At 4 weeks, we then added one small hole (1cm^2^) per side of the pot (the base already containing holes for drainage) for 2 groups of plants in 100ml pots. We then placed these plants, with their 100ml pot, into larger pots (500ml or 2000ml), which were filled with soil. Control plants were maintained in 100ml or 500ml pots as previously. Transferred plants (100→500ml and 100→2000ml) should encounter a similar level of mechanical stimulus to 100ml control plants, but have access to a much larger volume of soil – assuming they escape the original pot – and therefore a lower root density. In 100→500ml plants, peak tiller number in 11/12 cases was higher than in any of the 100ml control plants (Figure 7A), consistent with the idea that tillering is regulated by root density, not mechanical stimulus. However, production of tillers was slower than in 500ml control plants (Figure 7B). Furthermore, while 6/12 100→500ml plants produced tillers in the same range as 500ml control plants (9-19 tillers), the other 6/12 plants had reduced peak tiller number (4-8 tillers) relative to 500ml controls, although there was no significant difference overall between the groups (Figure 7A). In 100→2000ml plants, tiller production was also slower than in 500ml control pots (Figure 7B), and closely matched growth of 100→500ml plants in the two weeks after transfer. However, the peak tiller number in these plants eventually exceeded the 500ml control pots. In 7/12 plants, tiller numbers were higher than in any plant in the 500ml control pots, while in 5/12 plants, tiller numbers were in the same range as 500ml controls (9-19 tillers) (Figure 7A). In a single plant, a maximum of 7 tillers were produced, below the range of the 500ml control pots.

**Figure 7:**
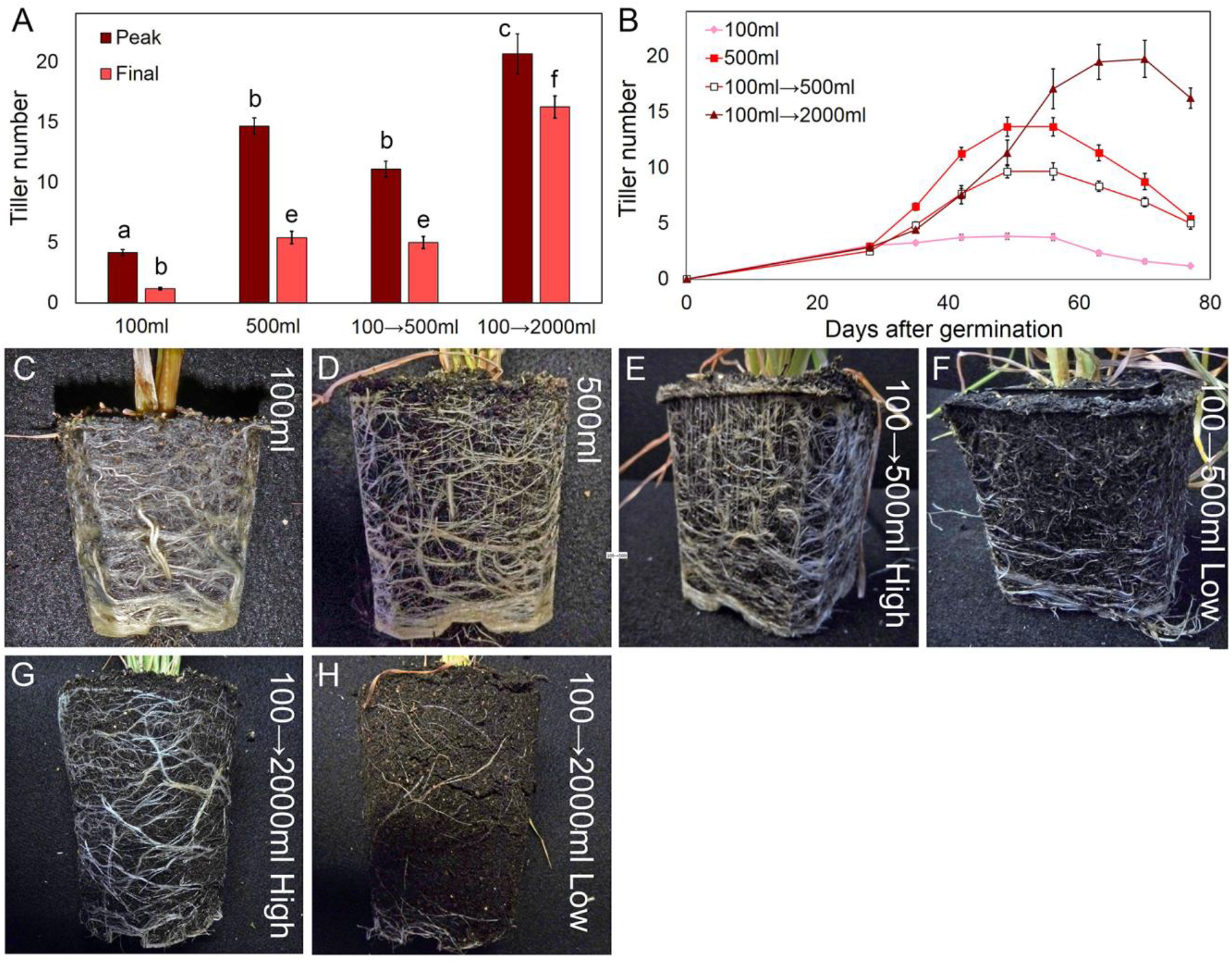
Shoot growth is correlated with effective root density. **A)** Graph showing peak and final tiller number in spring wheat grown in 100ml or 500ml pots, or grown in 100ml pots for 4 weeks then transferred (with pot) to 500ml or 2000ml pots. Error bars show s.e.m., n = 12. Bars with the same letter are not statistically different to each other (ANOVA+Tukey HSD). **B)** Graph showing mean tiller number in the 77 days after germination in spring wheat (Mulika) grown in 100ml or 500ml pots, or grown in 100ml pots for 4 weeks then transferred (with pot) to 500ml or 2000ml pots. Error bars indicate s.e.m, n=12. **C-H**) Root development visualised on the outer face of the soil volume in spring wheat (Mulika) grown in 100ml (C) or 500ml (D) pots, or grown in 100ml pots for 4 weeks then transferred (with pot) to 500ml (E, F) or 2000ml pots (G, H). E and G show high-tillering examples, F and H low-tillering examples.

The delay in tiller formation in 100→500ml and 100→2000ml plants is consistent with the plants temporarily remaining at high root density until the roots ‘escape’ into the outer soil jacket. Similarly, the low tiller values in certain transferred plants suggested that the roots of these plants may not have colonized the outer soil jacket effectively. To assess this idea, we examined root system growth in the atypically low-tillering 100→500ml and 100→2000ml plants, comparing them to high-tillering 100→500ml and 100→2000ml plants, and to 100ml and 500ml control plants, 6 weeks after transfer. In control plants, the soil was bound into a tight mass by extensive root growth, and could be removed from the pot and inspected externally (Figure 7C, D). Visible root density was similar in both pot sizes (Figure 7C, D). In high-tillering 100→500ml and 100→2000ml plants, the soil was also bound in a tight mass around the original pot, and visible root density was comparable to the control plants (Figure 7E, G). Conversely, in the low-tillering 100→500ml plants, the outer soil jacket was loosely bound, and in the 100→2000ml plants, completely unbound. Visible root density in the outer jacket was clearly much lower than in the controls and high-tillering transferred plants (Figure 7F, H). Thus, tillering in transferred plants was a function of the *utilised* soil volume. All transferred plants ultimately made roots in the outer soil jacket, but in lower-tillering plants this happened more slowly, and had a smaller effect on tillering during the window for tiller formation. When subsequently re-examined at 9 weeks after transfer, colonization of the outer root jacket was improved in these plants, but with little effect on tiller number.

Our results are thus consistent with tillering being determined by the effective root system density; that is to say, the mean density of roots in the utilised soil volume, rather than in the absolute soil volume. In 100ml pots, effective root system density rises more quickly than in 500ml pots, and tillering is inhibited more quickly in response. Plants in both pots ultimately cease tillering upon reaching a similar root system density. In transferred plants that efficiently colonize the outer soil jacket, effective root density increases more slowly than in 100ml control plants, allowing increased tillering, but in those plants that do not efficiently colonize the outer soil jacket, root system density still rises quickly, leading to partially inhibited tillering.

### Root system density connects volume restriction and crowding responses

As a final test of our root density hypothesis, we sought an alternative approach to increase root density without altering soil volume or mechanical impedance. To achieve this, we grew multiple plants of the same genotype in the same soil volume, reasoning that each individual plant would encounter increased root density much more quickly in this scenario, and respond by reducing its shoot growth. First, we grew either 1 or 5 spring wheat (Mulika) plants in 500ml of soil. Over a 15-week period, the growth of individual plants in the 5 per pot treatment was essentially identical to plants grown in 100ml of soil (in previous experiments)(Figure 8E). Thus, the early increase in root density when crowded by a factor of 5 makes the plants behave as if they were in in a soil volume 1/5^th^of the actual size. This strongly supports the idea that plants use root density to sense soil volume. Plants in the 1 per pot treatment finished their life with 3 or 4 tillers and ears, while each plant in the 5 per pot treatment finished with a single tiller and ear (Figure 8A, B). However, the total number of spikelets initiated per pot was practically identical between the treatments, as was the total shoot biomass per pot (Figure 8C, D). Thus collectively, the shoots in the 5 per pot treatment grew identically to the shoots in the 1 per pot treatment, showing remarkable cooperative behaviour. The variation between plants sharing a pot was small; the biomass is made up of 5 similar sized plants, rather than 1 or 2 dominant plants.

**Figure 8:**
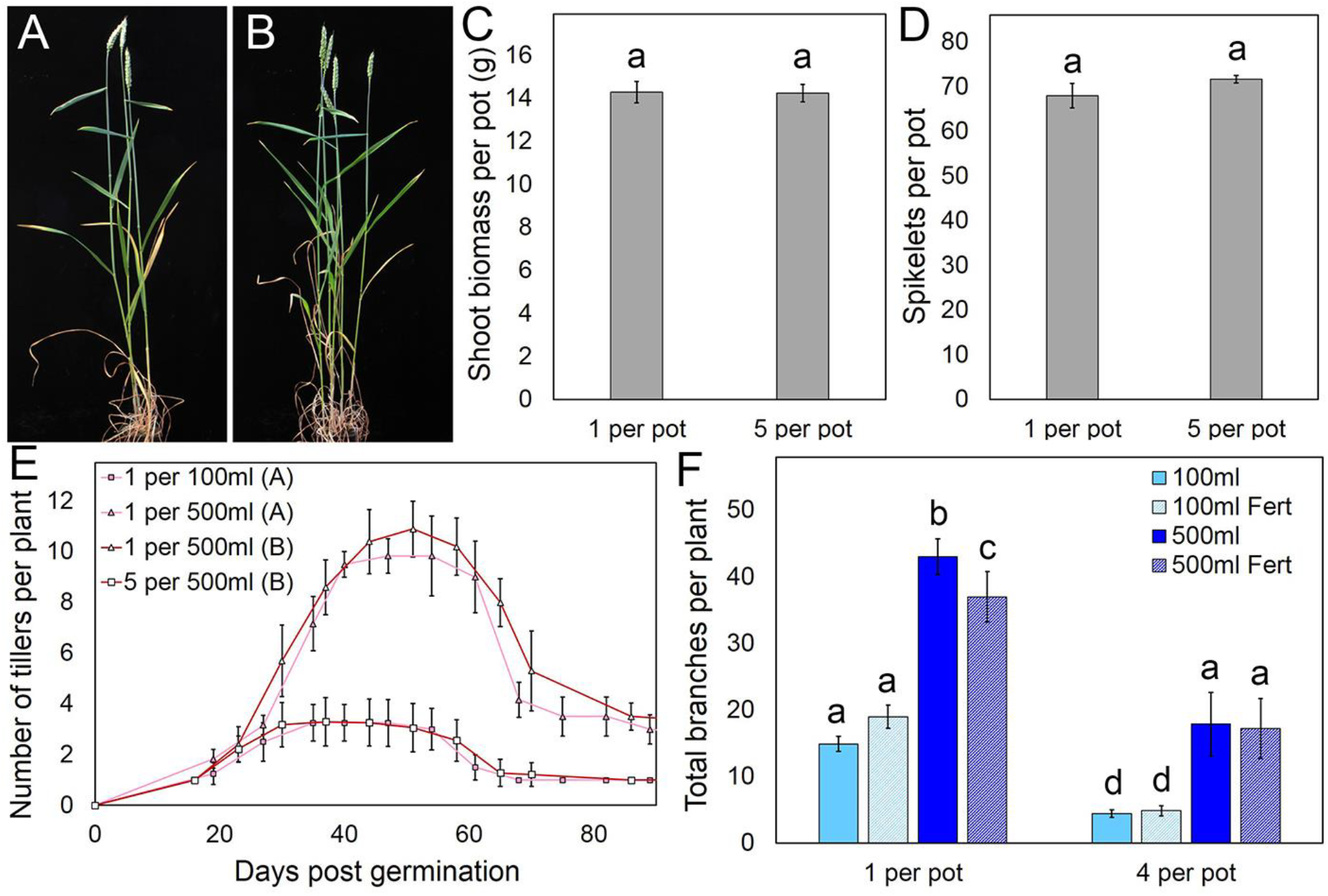
Root density determines shoot growth. **A,B)** Shoot development in spring wheat (Mulika) grown in 500ml of soil at a rate of 1 per pot (A) and 5 per pot (B). **C)** Graph showing total shoot biomass per pot in spring wheat (Mulika) grown in 500ml of soil at a rate of 1 per pot and 5 per pot. Error bars indicate s.e.m, n=10 pots. Bars with the same letter are not statistically different from each other (t-test). **D)** Graph showing total spikelets per pot in spring wheat (Mulika) grown in 500ml of soil at a rate of 1 per pot and 5 per pot. Error bars indicate s.e.m, n=10 pots. Bars with the same letter are not statistically different from each other (t-test). **E)** Graph showing mean tiller number in the 90 days after germination in spring wheat (Mulika) grown on 500ml of soil at a rate of 1 per pot and 5 per pot (red lines, dataset ‘B’). Data from Fig 4A showing single Mulika plants grown in 100ml and 500ml of soil (in a separate experiment) are superimposed (pink lines, dataset ‘A’). Error bars indicate s.e.m, n=6-12 for dataset A, n=10 for dataset B. **F)** Graph showing mean total branch number in wild-type (Col-0) Arabidopsis at end-of-life in plants grown on 100ml or 500ml of soil, treated with (‘Fert’) or without additional fertiliser, and at a rate of 1 or 4 plants per pot. Error bars indicate s.e.m, n=5-48. Bars with the same letter are not statistically different from each other (ANOVA+Tukey HSD).

To rule out possible confounding variables, we performed similar experiments using wild-type Arabidopsis, growing plants either 1 per pot or 4 per pot in 100ml or 500ml pots, thus varying root density by two different approaches. Within the 500ml pots, we crowded the plants into the same surface area as in the 100ml pots. This kept shading equal between the 4 per pot treatments in 100ml and 500ml, so changes in growth arose from changes in root density, rather than shading. In each treatment group, we also applied additional fertiliser to half the plants, to rule out nutrient deficiency as a causative agent. In plants grown at a rate of 1 per pot, the plants grew according to their soil volume, as anticipated, and were much larger in 500ml pots than 100ml pots (Figure 8F, Figure S3A). In both pot sizes, plants grown at a rate of 4 per pot were much smaller than the equivalent plants grown at a rate of 1 per pot (Figure 8F). However, the plants grown at 4 per pot in 500ml pots were much larger than those grown 4 per pot in 100ml pots, and indeed, were approximately the same size as those grown 1 per pot in 100ml (Figure 8F). There was little effect of additional fertiliser on the growth of any of these plants (Figure 8F). As with the wheat, the total shoot biomass in each pot size was similar whether 1 or 4 plants were grown in them, the plants again demonstrating highly cooperative growth. Our data from Arabidopsis and wheat therefore demonstrate that plants use the mean root density in their environment – irrespective of whether the roots are their own – to guide their shoot growth, and pro-actively avoid resource limitation. This allows them to sense their available soil volume, even in complex scenarios with multiple root systems present. These data serve as an excellent illustration of the remarkable ability of plants to sense their environment and to coordinate development over long distances in response.

## DISCUSSION

### The adaptive value of soil volume-sensing

Gardeners everywhere are familiar with concept of plants becoming ‘pot-bound’, and the need to transplant to larger pots. The restrictive growth effect of culturing plants in small pots has long been recognised, but has remained poorly characterised and understood (Poorter et al, 2012). There has been sufficient evidence to show that volume restriction is not a direct nutritional effect, but there has been neither a clear understanding of the ontogeny of root restrictive effects, nor their adaptive value (Poorter et al, 2012). Our data provide a framework for understanding the physiology and developmental biology of volume restriction. By sensing their own root density, plants can establish the volume of soil that they currently occupy. Coupled with detection of the concentration of nutrients in that soil volume, this would allow the plants to estimate the total nutrients likely to be available to them during their life cycle. Using this integrated value would be a more reliable guide for growth than relying on nutrient concentration alone. For instance, even if nutrient concentration is instantaneously high, if soil volume is low, the total available nutrients are low and the plant should restrict its growth to avoid resource limitation later in the life-cycle. It must be noted that in many of our experiments soil volume imposed a hard limit on growth, such that increasing nutrient concentration was not sufficient to increase growth. This may be because soil volume does not only predict nutrient availability, but also water availability. If the root system occupies a defined volume, this sets an upper limit on the availability of water, and therefore the maximum amount of shoot tissue that can be sustained, even in good conditions. The ability to pro-actively restrict shoot growth in response to soil volume therefore allows plants to complete their life-cycle almost irrespective of the starting resources.

An obvious question remains regarding the adaptive value of soil volume sensing, since plants do not encounter plant pots in the wild. There are several answers to this question. Firstly, plants may naturally encounter hard physical boundaries in several scenarios. For instance, in many places the soil layer is thin, and the hard, underlying bedrock may block the growth of root systems, causing volume restriction. Even in deep soil, larger species such as trees could become volume restricted in this way. In agricultural contexts, soil compaction caused by heavy machinery, or soil pans caused by ploughing may cause volume restriction, and thereby reduce yields. Secondly, our data show that no mechanical barrier is necessary for volume restriction to occur – it is the effective root density, rather than the soil volume *per se*, that causes volume restriction. Thus, even if there is no absolute barrier to root growth, if most of the root system is at high density, volume restriction would still occur. Thirdly, and perhaps most importantly, our data show that soil volume may be connected to the sensing of other plants in the rhizosphere.

### Soil volume and neighbour detection

The ability of plants to detect and respond to neighbouring plants through their root systems is well-established, although the literature surrounding this area is complex and sometimes confusing (Depuydt et al, 2014). Indeed, many studies have been criticised on the basis that soil volume is a confounding variable in their experimental design (Hess & De Kroon, 2007; Semchenko et al, 2008; Chen et al, 2012). Our results certainly support the importance of conducting experiments on plant neighbour detection in fixed soil volumes. Broadly speaking, older studies in this area deal with the generic ability of plants to detect their neighbours (usually unrelated) (e.g. Mahall & Callaway, 1990), and to discriminate their own roots from those of neighbouring plants (self/non-self-discrimination)(Falik et al, 2003; Gruntman & Novoplansky, 2004; Gersani et al, 2001). However, in this context self/non-self does not imply anything about relatedness, and indeed separated parts of the same original root system can be perceived as both self and non-self by a clonally derived plant (Gruntman & Novoplansky, 2004). More recently, studies have examined the ability of plants to differentially detect and respond to close kin, distant kin and non-kin (Dudley & File, 2007; Bierdrzycki et al, 2010; Crepy & Casal, 2016; Yang et al, 2018). Typically, cooperative behaviour is observed between close kin, with increasing competition as a function of genetic distance (Yang et al, 2018). However, the mechanism by which plants detect each other in the rhizosphere has remained obscure, with suggestions ranging from phytohormone signals, root exudates and electrical or mechanical signals (Depuydt et al, 2014). Most recently, jasmonic acid and (-)-loliolide have been proposed to function as exudates allowing neighbour detection, but simple chemicals such as these seem unlikely to encode sufficient information to allow discrimination of neighbours along a gradient of relatedness (Kong et al, 2018).

Our data strongly suggest that at least one of the mechanisms by which plants detect their neighbours is interchangeable with the mechanism by which they detect the density of their own root system, and hence their soil volume. In our crowding experiments, plants crowded by a factor of x behaved as if grown in a volume of soil 1/x^th^of the actual pot size, even if additional nutrients were provided. The most parsimonious explanation for this behaviour is that the plants detected root density in the pots indiscriminately, and restricted their shoot growth accordingly. Even though the density of their own roots was much lower than the total root density in the pot, they nevertheless became volume restricted. Thus, plants may detect each other in the rhizosphere at least partially through root density-sensing. It should be noted that the plants used for these experiments were completely inbred and ∼genetically identical in the case of Arabidopsis, and in wheat to a slightly lesser degree. Thus, in effect, the plants were detecting the density of their ‘own’ roots, perhaps explaining the extremely cooperative outcome in these experiments. Despite the closeness of their genetic relationship, the ability of these plants to precisely detect and respond to each other, and the similarity of this response to their response to soil volume is remarkable. In line with other recent studies, we suggest that an exudate-based system most likely accounts for these remarkable abilities. We therefore suggest a model in which plants exude and detect a chemical across their root system that allows them to both perceive the density of their own roots together with the roots of any close kin, with the adaptive value of both neighbour/kin-recognition and soil volume-sensing (Figure 9). This model cannot explain the competitive responses that occur between non-kin plants, so we suggest that ‘self/kin-perception’ through this root density-sensing system might downregulate competitive responses that are otherwise triggered by detection of ‘generic’ neighbour exudates such as jasmonate and (-)-loliolide.

**Figure 9:**
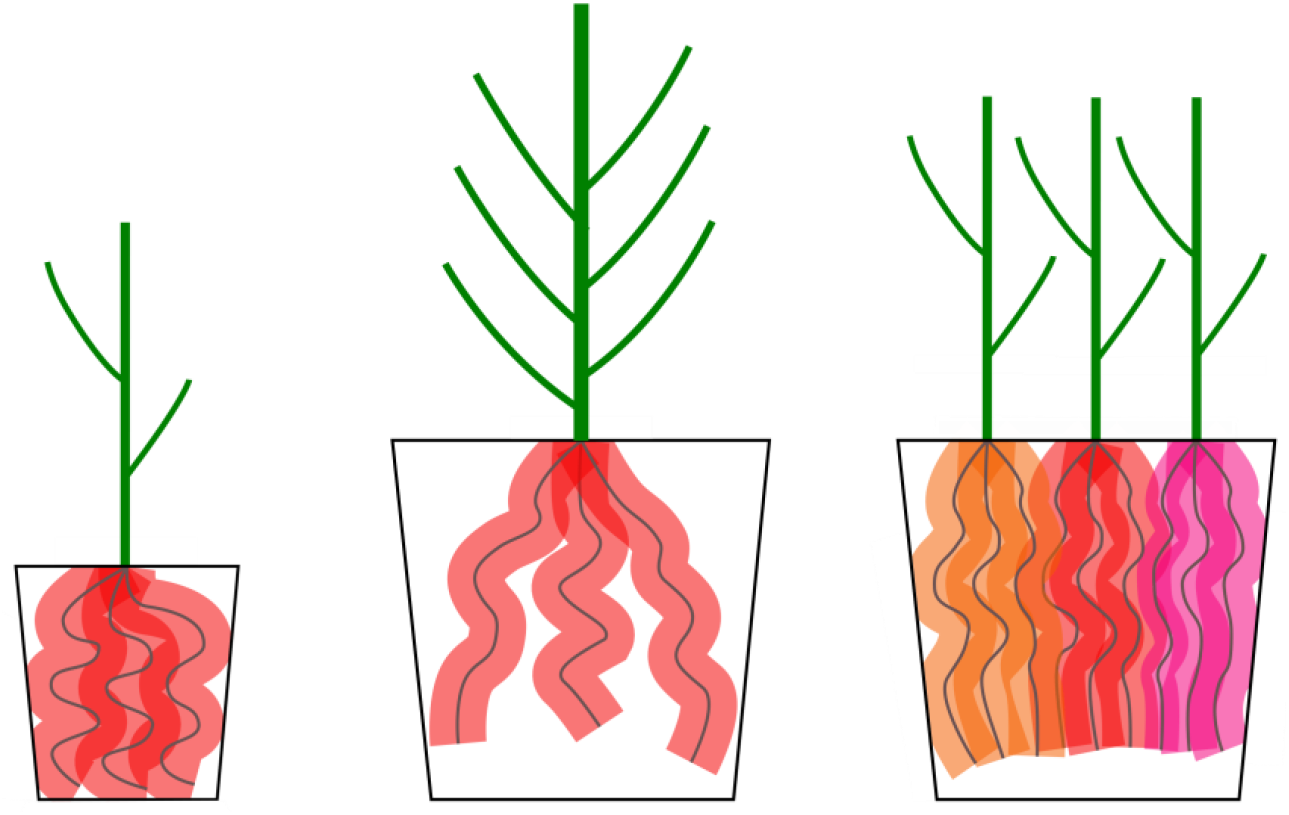
A model for soil volume sensing and kin recognition. Early in development, each plant has produced the same number and length of roots (thin grey lines), and has exuded the same amount of compounds into the rhizosphere (shaded areas around the roots). In plants grown in small pots (left) or at high density (right), the density of exudates rises much more quickly, alerting plants to possible resource limitations. Root-shoot signalling is modified to reduce branching and shoot biomass accumulation *before* root growth stops, and *before* resources are exhausted. In plants grown in large pots (centre), root density does not hit a critical level until later in development, and more shoot branches and biomass are formed.

### Root density, nutrient use efficiency and yield

Nutrient use efficiency (NUE) is a persistent problem in crop plants, with typically only 50% of nutrients available to the plants taken up (Fageria and Baligar, 2005; Raun and Johnson, 1999). Fertiliser applications make up a major part of the economic and environmental costs of arable agriculture, and if NUE could be increased, fertiliser applications could be correspondingly reduced with all-round benefits (Garnett, et al, 2009; Dobermann and Cassman, 2005). Our data show that plants do not respond to additional nutrient availability under conditions of high root density (volume restriction or crowding); they become so ‘cautious’ about the potential availability of resources later in the life cycle that they ignore the extra nutrients. Since most crops are grown at high density, and often in shallow or compacted soil, they are likely to experience very high root densities, and corresponding decreases in NUE, as a simple structural response to their environment.

At a broader scale, our data show that shoot biomass, branching and yield can be limited even in the presence of sufficient nutrients, due to other environmental cues. This is true even for crop plants like wheat and barley that have been bred for thousands of years to grow at high density and for high yields. Mathematical modelling of crop growth suggests that typical crop yields fall well short of the potential yields that should be possible with available resources (Schils et al. 2018; Foulkes et al. 2011). For instance, in the UK, average winter wheat yields are around 9 tonnes/hectare, compared to a theoretical potential of 22 tonnes/hectare (Sylvester-Bradley and Wiseman, 2004). We believe that the inherent caution of plants with respect to future resource availability acts as a major structural limitation on yield in all crop species, and that creating less cautious crops could produce step-changes in yields without altering inputs. Our data suggest that reducing the sensitivity of crops to their root density could be one relatively straightforward approach to achieving this goal.

## MATERIALS & METHODS

### Plant growth conditions

Arabidopsis plants for phenotypic and microsurgical experiments were grown on Petersfield No. 2 compost, or a sand/vermiculite mix, under a standard 16 h/8 h light/dark cycle (20°C/16°C), primarily in controlled environment rooms with light provided by white fluorescent tubes at intensities of ∼120μmol/m^2^s^−1^.

Wheat, barley and oilseed rape were grown on Petersfield No. 2 compost, or sand/vermiculite (50:50), sand/perlite (50:50) or vermiculite, in greenhouses with supplemental LED (wheat, barley) or sodium lamps (oilseed rape) to an average intensity of ∼250μmol/m^2^s^−1^. The Arabidopsis experiment shown in Figure 3 was grown under comparable conditions.

### Plant materials

The lines used in this study were Arabidopsis wild-type Col-0, spring wheat varieties Mulika and Willow, spring barley varieties Propino and Charon and spring oilseed rape variety Heros.

### Fertiliser treatments

We used Arabidopsis Thaliana Salts (ATS) (Wilson et al, 1990) as a standard modular fertiliser, and we varied the nitrate concentration by replacing nitrate ions with chloride. Standard N fertiliser was 0.015M nitrate, low N fertiliser was 0.0015M nitrate. Plants grown on sand and vermiculite received 5ml of ATS + 5ml water once per week in place of watering. Plants grown on compost received 5ml of standard ATS + 5ml of water (Arabidopsis) or 10ml of standard ATS (wheat, barley, OSR) every week in place of watering.

### Phenotypic assessments

Biomass measurements were made at the end of life unless stated otherwise. For the data in Figure 2A, Figure 5, 4 plants were destructively sampled for biomass at 2, 4 and 6 weeks after germination. For wheat, barley and Arabidopsis, we measured dry biomass and for oilseed rape, fresh biomass. For some experiments, we separated wheat biomass into straw and ear biomasses. Where measured, seed biomass was assessed by threshing ears, and sieving off chaff.

Branching in Arabidopsis and oilseed rape was assessed at the end of life (∼7 or ∼16 weeks respectively). Branches of all classes (primary, secondary, tertiary, quaternary) were counted separately and combined for total branch numbers. Tillering in wheat in barley was assessed every week from 3 weeks after germination. Ear, spikelet and seed counts were made at the end of life (∼15 weeks after germination).

Visual assessments of root growth were performed as described in the text. For the biomass data in Figure 5, roots were extracted from vermiculite and washed to remove any remaining particles, dried and then measured.

## ACKNOWLEDGEMENTS

This work was supported by funding from the N8 Agrifood program. We thank Stefan Vanneste for helpful suggestions, and Besiana Sinanaj for technical assistance.

## AUTHOR CONTRIBUTIONS

CDW, CHW, TB performed experiments and analysed the data. CHW & TB designed the study. All authors contributed to writing the manuscript.

---

**Figure S1:**
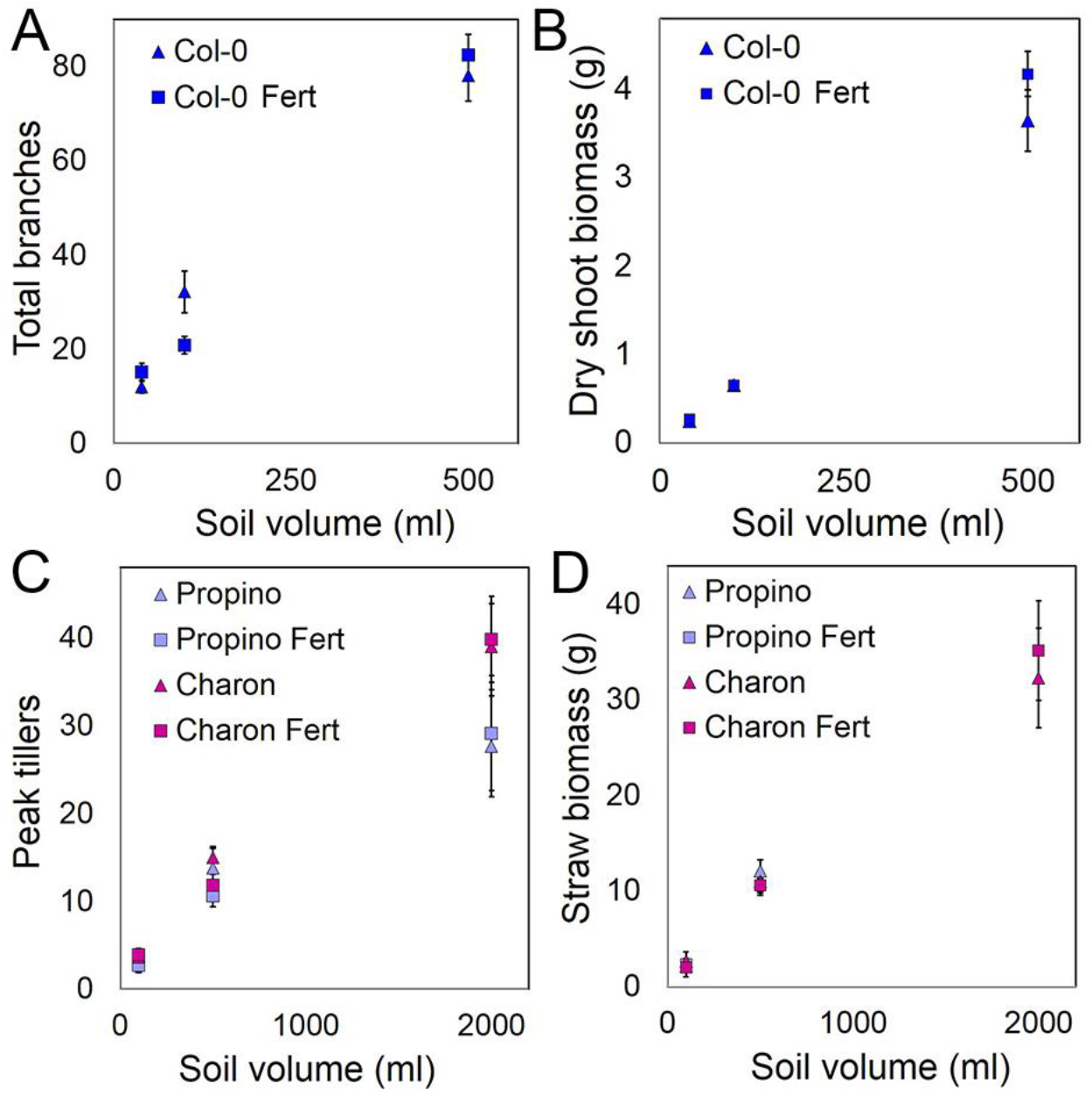
Soil volume directly influences plant growth. **A,B)** Graphs showing the relationship between soil volume and mean total branch number (A) and mean dry shoot biomass in grams (B) in Arabidopsis grown in 50, 100, and 500ml of soil, with (‘Fert’) or without additional fertiliser. Error bars indicate s.e.m, n=10-12. **C,D)** Graphs showing the relationship between soil volume and mean peak tiller number (C), mean dry straw biomass in grams (D), in two varieties of spring barley (Charon and Propino) in 100, 500 and 2000ml of soil, with (‘Fert’) or without additional fertiliser. Error bars indicate s.e.m, n=6-12.

**Table S1:** Soil volume directly influences wheat reproductive architecture. Table showing reproductive parameters in spring wheat (Mulika and Willow) grown in 100, 500 and 2000ml pots, with or without additional fertiliser (‘Fert’). Data are means ± s.e.m., n=6-12.

| Variety | Volume (ml) | Fert | Ear |  | Spikelet | Seed |  |
| --- | --- | --- | --- | --- | --- | --- | --- |
|  |  |  | Number | Mass (g) | Number | Number | Mass (g) |
| Mulika | 100 | - | 1.000<br>±0.000 | 1.089<br>±0.117 | 22.500<br>±0.506 | 26.545<br>±2.934 | 0.869<br>±0.123 |
|  | 100 | + | 0.917<br>±0.083 | 0.815<br>±0.102 | 21.273<br>±0.367 | 22.000<br>±2.111 | 0.551<br>±0.091 |
|  | 500 | - | 2.833<br>±0.152 | 6.882<br>±0.411 | 68.333<br>±3.896 | 113.167<br>±21.433 | 5.232<br>±0.343 |
|  | 500 | + | 3.000<br>±0.236 | 6.668<br>±0.513 | 70.167<br>±5.474 | 140.167<br>±11.680 | 5.089<br>±0.465 |
|  | 2000 | - | 9.333<br>±0.733 | 21.548<br>±1.716 | 231.333<br>±21.863 | 452.167<br>±39.236 | 14.903<br>±1.169 |
|  | 2000 | + | 9.333<br>±0.561 | 17.657<br>±2.356 | 214.833<br>±13.016 | 360.167<br>±30.994 | 14.402<br>±2.519 |
| Willow | 100 | - | 1.000<br>± 0.123 | 1.474<br>± 0.071 | 20.455<br>± 1.976 | 35.727<br>± 2.548 | 1.119<br>± 0.047 |
|  | 100 | + | 0.667<br>± 0 | 0.460<br>± 0.088 | 7.727<br>± 3.297 | 5.286<br>± 1.633 | 0.170<br>± 0.057 |
|  | 500 | - | 4.500<br>± 0.204 | 6.658<br>± 0.420 | 92.500<br>± 7.623 | 145.666<br>± 16.062 | 5.056<br>± 0.607 |
|  | 500 | + | 4.333<br>± 0.304 | 6.590<br>± 0.455 | 82.000<br>± 6.743 | 138.000<br>± 8.307 | 4.951<br>± 0.424 |
|  | 2000 | - | 7.833<br>± 2.522 | 16.592<br>± 4.449 | 210.500<br>± 45.596 | 307.000<br>± 86.420 | 10.275<br>± 2.947 |
|  | 2000 | + | 11.333<br>± 1.981 | 17.395<br>± 4.010 | 279.000<br>± 30.640 | 359.833<br>± 57.742 | 11.126<br>± 2.107 |

## REFERENCES

Bar-Tal A, Feigin A, Sheinfeld S, Rosenberg R, Sternbaum B, Rylski I, Pressman E. (1995) Root restriction and N-NO3 solution concentration effects on nutrient uptake, transpiration and dry matter production of tomato. Sci Hortic 63, 195–208

Bar-Tal A, Pressman E (1996) Root Restriction and Potassium and Calcium Solution Concentrations Affect Dry-matter Production, Cation Uptake, and Blossom-end Rot in Greenhouse Tomato. J Amer Soc Hort Sci 121, 649–655

Bennett T, Leyser O (2014). The auxin question: a philosophical overview. In: Auxin and its Role in Plant Development (Eds: Zazimalova E, Petrasek J, Benkova E). Springer, Berlin. 3–19.

Biedrzycki ML, Jilany TA, Dudley SA, Bais HP (2010). Root exudates mediate kin recognition in plants. Commun Integr Biol 3, 28–35.

Carmi, A. and Heuer, B. (1981). The Role of Roots in Control of Bean Shoot Growth. Ann Bot 48 519–527.

Chen BJ, During HJ, Anten NP (2012). Detect thy neighbor: identity recognition at the root level in plants. Plant Sci 95, 157–166.

Crepy MA, Casal JJ (2016). Kin recognition by self-referent phenotype matching in plants. New Phytol 20, 15–16.

de Jong M, George G, Ongaro V, Williamson L, Willetts B, Ljung K, McCulloch H, Leyser O (2014). Auxin and strigolactone signaling are required for modulation of Arabidopsis shoot branching by nitrogen supply. Plant Physiol. 166, 384–95.

Depuydt S (2014). Arguments for and against self and non-self root recognition in plants. Front Plant Sci 5:614

Dobermann A, Cassman KG (2005). Cereals and nitrogen use efficiency are drivers of future nitrogen fertilizer consumption. Science in China Series C: Life Sciences 48, 745–758.

Dudley SA, File AL (2007). Kin recognition in an annual plant. Biol Lett. 3, 435–438.

Fageria NK and Baligar VC (2005). Enhancing Nitrogen Use Efficiency in crop plants. Advances in Agronomy, 88 97–105.

Falik O, Reides P, Gersani M, Novoplansky A (2003) Self/non-self discrimination in roots. J Ecol 91, 525–531.

Foulkes MJ, Slafer GA, Davies WJ, Berry PM, Sylvester-Bradley R, Martre P, Calderini DF, Griffiths S, Reynolds MP (2011). Raising yield potential of wheat. III. Optimising partitioning to grain while maintaing lodging resistance. J Exp Bot 62, 469–486.

Garnett T, Conn V, Kaiser BN (2009). Root based approaches to improving nitrogen use efficiency in plants. Plant Cell Environ 32, 1272–1283.

Gersani M, Brown JS, O’Brien EE, Maina GM, Abramsky Z (2001). Tragedy of the commons as a result of root competition. Journal of Ecology 89, 660–669.

Gruntman M, Novoplansky A (2004). Physiologically mediated self/non-self discrimination in roots PNAS USA 101, 3863–3867.

Hameed MA, Reid JB, Rowe RN (1987). Root Confinement and its Effects on the Water Relations, Growth and Assimilate Partitioning of Tomato (Lycopersicon esculentum Mill). Ann Bot 59, 685–692.

Hess L, De Kroon, H. (2007). Effects of rooting volume and nutrient availability as alternative explanation for root self/non-self discrimination. J Ecol 95, 241–251.

Ismail MR, Davies WJ (1998), Root restriction affects leaf growth and stomatal response: the role of xylem sap ABA. Sci Hortic 74, 257–268.

Kharkina TG, Ottosen CO, Rosenqvist E (1999). Effects of root restriction on the growth and physiology of cucumber plants. Physiol Plant 105, 434–441.

Kohlen W, Charnikhova T, Liu Q, Bours R, Domagalska MA, Beguerie S, Verstappen F, Leyser O, Bouwmeester H, Ruyter-Spira C (2011). Strigolactones are transported through the xylem and play a key role in shoot architectural response to phosphate deficiency in nonarbuscular mycorrhizal host Arabidopsis. Plant Physiol 155, 974–987.

Krizek DT, Carmi A, Mirecki RM, Snyder FW, Bunce JA (1985) Comparative Effects of Soil Moisture Stress and Restricted Root Zone Volume on Morphogenetic and Physiological Responses of Soybean [Glycine max (L.) Merr.]. J Exp Bot 36, 25–38.

Mahall B, Callaway RM (1991). Root communication among desert shrubs. PNAS USA 88, 874–876.

Matsubayashi Y (2014). Post-translationally modified small-peptide signals in plants. Annu Rev Plant Biol. 65, 385–413.

Müller D, Waldie T, Miyawaki K, To JP, Melnyk CW, Kieber JJ, Kakimoto T, Leyser O (2015). Cytokinin is required for escape but not release from auxin mediated apical dominance. Plant J 82, 874–86.

Murphy GP, Van Acker R, Rajcan I, Swanton CJ (2017). Identity recognition in response to different levels of genetic relatedness in commercial soya bean. R Soc Open Sci. 4, 160879.

Osakabe Y, Osakabe K, Shinozaki K, Tran LS. (2014). Response of plants to water stress. Front Plant Sci 5:86.

Raun WR, Johnson GV (1999). Improving nitrogen use efficiency for cereal production. Agronomy J 91, 357–363.

Schils R, Olesen JE, Kersebaum KC, Rijk B, Oberforster M, Kalyada V, et al. (2018). Cereal yield gaps across Europe. Eur J Agronomy 101, 109–120.

Semchenko M, Hutchings MJ, John EA (2007). Challenging the tragedy of the commons in root competition: confounding effects of neighbour presence and substrate volume. J Ecol 95, 252–260.

Shemesh H, Zaitchik B, Acuña T, Novoplansky A. (2012). Architectural plasticity in a Mediterranean winter annual. Plant Signal Behav 7, 492–501.

Shi K, Ding XT, Dong DK, Zhou YH, Yu JQ (2008) Root restriction-induced limitation to photosynthesis in tomato (Lycopersicon esculentum Mill.) leaves. Sci Hortic 117, 197–202

Sylvester-Bradley R, Wiseman J (2005). Yields of Farmed Species: constraints and opportunities in the 21st century. Proceedings of University of Nottingham Easter School Series, June 2004, Sutton Bonington, UK. 2005. pp. xii + 651pp.

Ternesi M, Andrade AP, Jorrin J, Benlloch M (1994). Root-shoot signalling in sunflower plants with confined root systems. Plant Soil 166, 31–36

Umehara M, Hanada A, Yoshida S, Akiyama K, Arite T, Takeda-Kamiya N, Magome H, Kamiya Y, Shirasu K, Yoneyama K, Kyozuka J, Yamaguchi S (2008). Inhibition of shoot branching by new terpenoid plant hormones. Nature 455, 195–200.

Waldie T, Leyser O (2018). Cytokinin Targets Auxin Transport to Promote Shoot Branching. Plant Physiol. 177, 803–818.

Walker C, Bennett T (2018). Forbidden Fruit: dominance relationships in the control of shoot architecture. Annual Plant Reviews Online, apr0640.

Wilson AK, Pickett FB, Turner JC, Estelle M (1990). A dominant mutation in Arabidopsis confers resistance to auxin, ethylene and abscisic acid. Mol Gen Genet. 222, 377–383.

Yang XF, Li LL, Kong CH (2018). Kin recognition in rice (Oryza sativa) lines. New Phytol 220, 567–578.

Yong JWH, Letham DS, Wong SC, Farquhar GD (2010) Effects of root restriction on growth and associated cytokinin levels in cotton (Gossypium hirsutum). Funct. Plant Biol. 37, 974–984.

